# Vascular Microbleeds Without Brain Atrophy: A Microvascular Signature of Mid-Stage 5xFAD Pathology

**DOI:** 10.64898/2025.12.08.692809

**Authors:** Xiuli Yang, Yuguo Li, Adnan Bibic, Jiekang Wang, Mei Wan, Wenzhen Duan, Hanzhang Lu, Zhiliang Wei

## Abstract

Cerebral microbleeds are increasingly recognized as a downstream manifestation of vascular injury in Alzheimer’s disease (AD), arising secondary to cerebral amyloid angiopathy (CAA). Here, we examined the pathological specificity of microbleeds by comparing an amyloidosis mouse model (5xFAD) with a small-vessel disease (SVD) model characterized by vascular smooth-muscle cell loss. *In vivo* multimodal MRI, including gradient-echo, spin-echo, and diffusion-weighted imaging, was complemented by *ex vivo* high-resolution anatomical scans for validation. Both *in vivo* and *ex vivo* gradient-echo MRI consistently revealed hippocampal microbleeds in the 5xFAD model without macroscopic atrophy or ventricular enlargement, whereas no microbleeds or blood-brain barrier disruption were detected in the SVD model. Diffusion-weighted MRI further showed region-specific alterations in apparent diffusion coefficient within the midbrain of 5xFAD mice, but not in other regions or in the SVD cohort. These findings indicate that microbleeds are a pathology-specific marker of amyloid-related vascular injury. The imaging evidence underscores the potential of microbleeds as a disease-specific biomarker for detecting amyloid-driven vascular fragility and refining diagnostic and therapeutic strategies for AD.

## Introduction

Microbleeds are increasingly recognized as a neuroimaging hallmark of cerebrovascular injury and can be sensitively detected using T_2_*-weighted or susceptibility-weighted MRI.^1,2^ In Alzheimer’s disease (AD), microbleeds exhibit a predominantly lobar distribution and are strongly associated with cerebral amyloid angiopathy (CAA), which arises from amyloid-β deposition within cortical and leptomeningeal vessel walls.^3,4^ In patients with AD, microbleed burden correlates with accelerated cognitive decline, disruption of neurovascular integrity, and increased risk of spontaneous intracerebral hemorrhage,^5,6^ underscoring their role as an imaging indicator of amyloid-associated vascular fragility. Furthermore, microbleeds frequently co-occur with parenchymal amyloid deposition,^7^ neuroinflammation,^8^ ventricular enlargement,^9^ white matter hyperintensity,^10^ and blood-brain barrier dysfunction,^11^ linking them to broader neurovascular pathology rather than isolated hemorrhagic foci.

Cerebral small vessel disease (SVD) represents a major contributor to dementia syndromes^12^ and encompasses diverse non-amyloid vasculopathies, including hypertensive arteriopathy,^13^ cerebral autosomal dominant arteriopathy with subcortical infarcts and leukoencephalopathy (CADASIL),^14^ and other inherited or sporadic vascular pathologies.^15^ While cerebral microbleeds are also observed in SVD, their spatial distribution, burden, and underlying vascular drivers differ fundamentally from those seen in AD.^16,17^ SVD-associated microbleeds most frequently localize to deep brain regions, including the basal ganglia, thalamus, and brainstem.^1^ These divergent anatomical and biological patterns suggest that microbleeds are not a uniform consequence of small vessel injury, but rather encode disease-specific vascular pathophysiology.

In the literature, a diverse range of animal models has been developed to study AD and SVD pathologies.^18–20^ The 5xFAD mouse model co-expresses five familial AD mutations in *APP* and *PSEN1*, resulting in rapid accumulation of parenchymal amyloid-β plaques, progressive gliosis, neurodegeneration, and neurovascular dysfunction.^21^ In contrast, CADASIL is an inherited SVD caused by *NOTCH3* mutations and is defined by vascular smooth muscle cell degeneration, arteriopathy, and chronic small vessel injury without amyloid pathology.^22^ To determine whether cerebral microbleeds reflect a pathology-specific consequence of amyloid-associated vascular injury rather than a general outcome of small-vessel impairment, we performed a side-by-side MRI investigation in 5xFAD and CADASIL mice. 5xFAD mice were evaluated at 9–12 months, an age range characterized by established amyloid deposition^21^ and emerging vascular injury, with microbleeds verified using histology and *ex vivo* MRI. CADASIL mice, modeling progressive non-amyloid arteriopathy, were examined at 9, 13, and 20 months to capture potential age-dependent vascular changes.

## Materials and Methods

### General procedures

Experimental protocols for this study were approved by the Johns Hopkins Medical Institution Animal Care and Use Committee and conducted in accordance with the National Institutes of Health guidelines for the care and use of laboratory animals. Data reporting complied with the ARRIVE 2.0 guidelines. All procedures were carefully designed to minimize discomfort and stress to the animals. Mice were housed in a quiet environment under a 12-h light/dark cycle with *ad libitum* access to food and water. A total of 81 mice (age: 9–20 months; body weight: 20-58 grams; 42 females [F], 39 males [M]) were investigated in this study. An amyloidosis model 5xFAD^21^ (age: 9-12 months; *n* = 28; 16F12M) were examined with multimodal structural MRI techniques. In a subgroup of the 5xFAD model (WT: *n* = 5, 1F4M; 5xFAD: *n* = 5, 3F2M), mouse brains were extracted for *ex vivo* MRI scans. A CADASIL model^22^ (age: 9, 13, and 20 months; *n* = 53; 26F27M), a transgenic form of SVD, was included for verifying the pathological specificity of microstructural changes. Information of mouse numbers was detailed in Table 1. For mice scanned on the same day, the experimental order was randomized according to a previously reported scheme^23^.

**Table 1.**
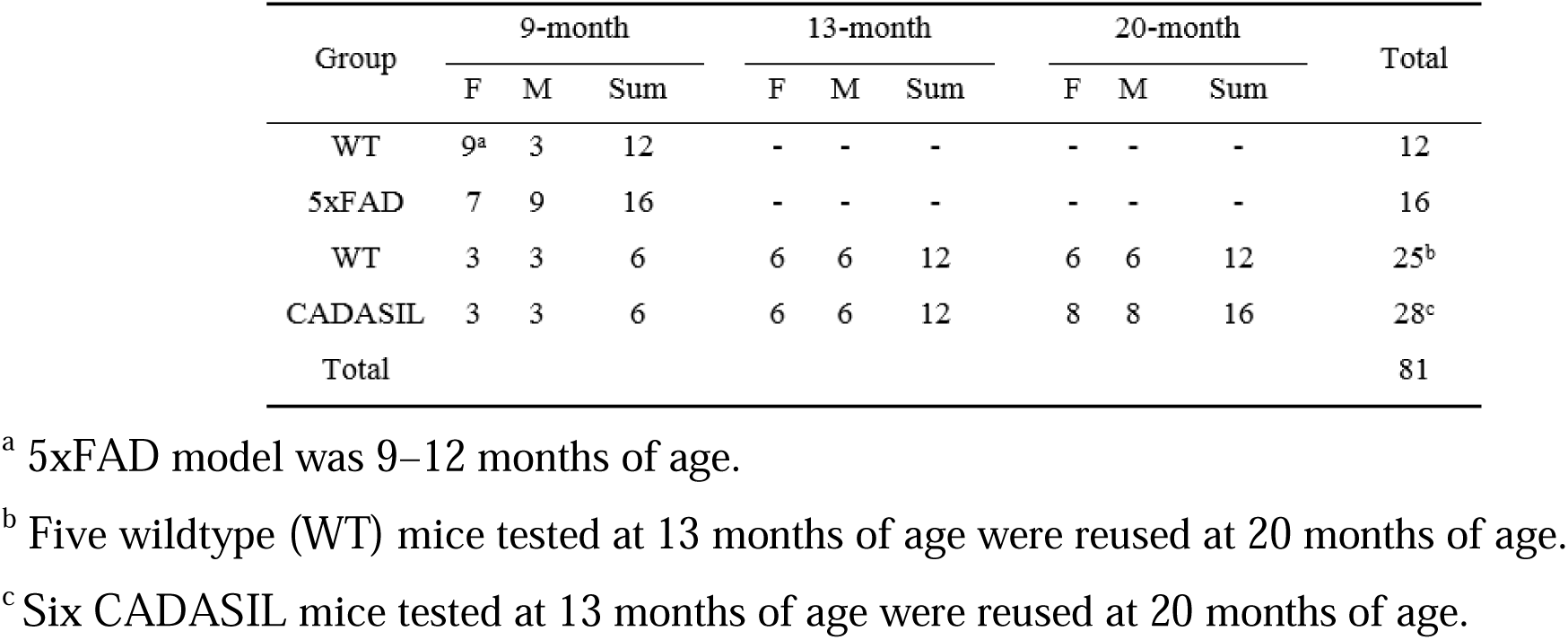
Mouse numbers for each experiment across the examined time points.

| Group | 9-month |  |  | 13-month |  |  | 20-month |  |  | Total |
| --- | --- | --- | --- | --- | --- | --- | --- | --- | --- | --- |
|  | F | M | Sum | F | M | Sum | F | M | Sum |  |
| WT | 9<sup>a</sup> | 3 | 12 | - | - | - | - | - | - | 12 |
| 5xFAD | 7 | 9 | 16 | - | - | - | - | - | - | 16 |
| WT | 3 | 3 | 6 | 6 | 6 | 12 | 6 | 6 | 12 | 25<sup>b</sup> |
| CADASIL | 3 | 3 | 6 | 6 | 6 | 12 | 8 | 8 | 16 | 28<sup>c</sup> |
| Total |  |  |  |  |  |  |  |  |  | 81 |

<sup>a</sup> 5xFAD model was 9–12 months of age.
<sup>b</sup> Five wildtype (WT) mice tested at 13 months of age were reused at 20 months of age.
<sup>c</sup> Six CADASIL mice tested at 13 months of age were reused at 20 months of age.

### MRI

An 11.7 T Bruker Biospec system (Bruker, Ettlingen, Germany) with a horizontal bore and actively shielded pulsed field gradient (maximal intensity: 0.74 T/m) was used for imaging. Data were acquired using a 72-mm quadrature volume resonator as the transmitter and a four-element (2 × 2) phased-array coil as the receiver. B0 homogeneity across the mouse brain was optimized using global shimming (up to the second order) based on a subject-specific pre-acquired field map. Inhalational isoflurane was used as the anesthetic following a previously reported scheme ^24,25^. Respiration was observed during the experiments using an MRI-compatible monitoring system (SA Instruments, Stony Brook, USA). Isoflurane dose was adjusted as needed to maintain a respiration rate of 70–120 breaths per minute. Experiments were terminated if respiration rates dropped below 50 breaths per minute for more than two minutes. A temperature-controlled water circulation system embedded in the scanner was used to maintain body temperature in the mice.

Figure 1 illustrates the experimental design of this study. Gradient-echo (GRE) MRI was performed to assess microbleeds, while spin-echo (SE) MRI was used to evaluate hyperintensities, parenchymal tissue volume, and cerebrospinal fluid (CSF) volume. Diffusion-weighted imaging (DWI) was applied to examine microstructural tissue properties through the apparent diffusion coefficient (ADC). The 5xFAD model served as the primary focus of this study, with CADASIL mice included as a comparative group to identify pathology-specific alterations.

**Figure 1.**
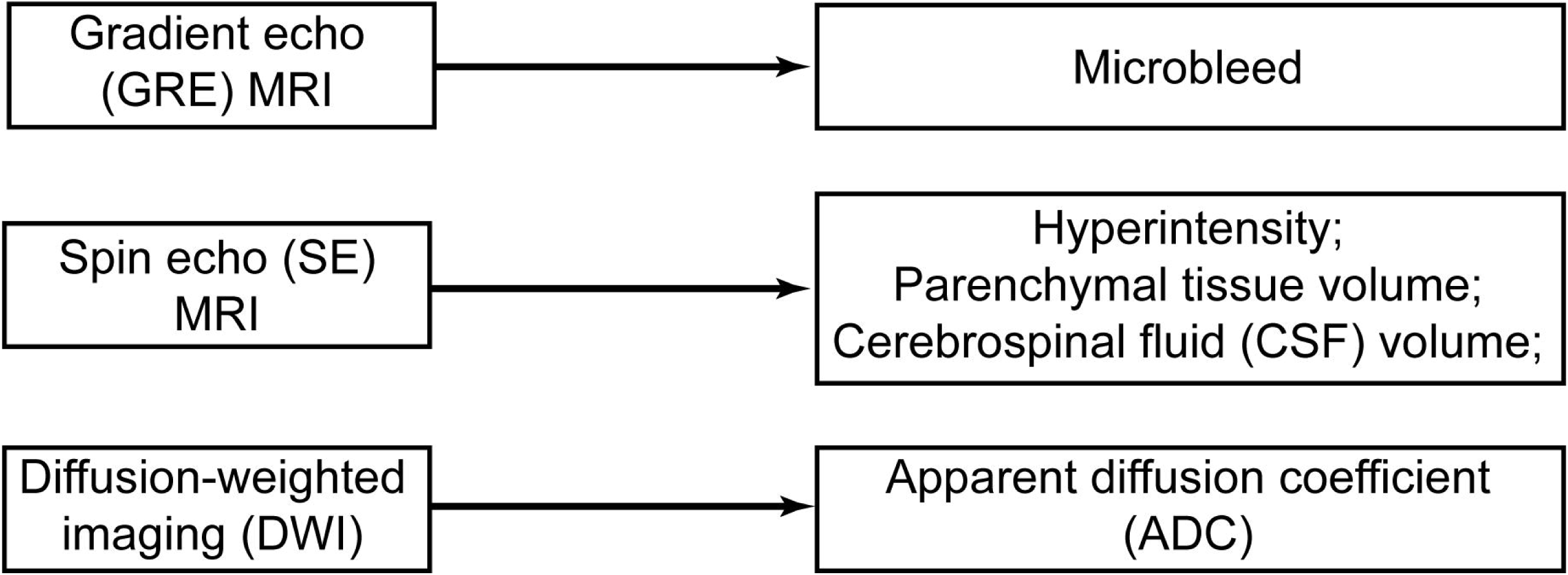
Schematic diagram of the study design.

Key acquisition parameters for the GRE MRI in the 5xFAD model were as follows: TR = 250 ms; TE = 12.0 ms; field of view = 15 (left–right) × 7.5 (ventral–dorsal) × 15 (rostral–caudal) mm^3^; matrix size = 256 × 128 × 48; spatial resolution = 59 × 59 × 313 µm^3^; receiver bandwidth = 110 kHz; partial-Fourier factor = 0.75 along the phase-encoding direction; and scan duration = 19.2 min with 3D acquisition. For the extracted brains of 5xFAD model, the *ex vivo* GRE scan followed the *in vivo* parameters except that: field of view = 12 (left–right) × 7.5 (ventral–dorsal) × 15 (rostral–caudal) mm^3^; matrix size = 256 × 128 × 256; spatial resolution = 47 × 59 × 59 µm^3^; and scan duration = 102.4 min. In the CADASIL model, key acquisition parameters for GRE MRI were: TR = 900/700/900 (9/13/20 months of age, respectively), TE = 10.0 ms; field of view = 15 × 15 mm^2^; matrix size = 256 × 256; spatial resolution = 59 × 59 µm^2^, slice thickness = 0.75 mm; 16 axial slices; average number = 4; partial-Fourier factor = 0.75/0.67/0.75 (9/13/20 months of age) along the phase-encoding direction; and scan duration = 11.5/8.0/11.5 min. In a subgroup of CADASIL model at 20 months of age (*n* = 4 [2F2M] WT, *n* = 8 [4F4M] CADASIL), the *in vivo* GRE protocol for the 5xFAD model was used.

Our previous report revealed altered permeability-surface area product (PS), a quantitative marker for BBB integrity, in the 5xFAD mice from 3 months of age.^26^ Therefore, we did not repeat this measurement for 5xFAD mice. In the CADASIL mice, Water-extraction-with-phase-contrast-arterial-spin-tagging (WEPCAST) MRI^27,28^ was applied to measure PS following the Renkin-Crone equation,^29,30^ i.e., *PS* = -ln (1 - *E*) *CBF* , where E and CBF denote water extraction fraction and cerebral blood flow, respectively. Key acquisition parameters of WEPCAST MRI were^28^: repetition time (TR) = 3000 ms; echo time (TE) = 11.8 ms; field of view = 15 × 15 mm^2^; matrix size = 96 ×96; slice thickness = 1.0 mm; labeling duration = 1200 ms; inter-labeling-pulse delay = 1.0 ms; post-labeling delay = 100 ms; partial Fourier factor = 0.67 along the phase-encoding direction; receiver bandwidth = 300 kHz; and scan duration = 4.0 min with a two-segment echo-planar-imaging (EPI) acquisition covering the midsagittal plane of brain. In addition, a reported phase-contrast protocol^24^ was used to estimate CBF.

Key imaging parameters for the SE sequence in the 5xFAD model were as follows: TR/TE = 1500/5.4 ms; field of view = 15 (left–right) × 7.5 (ventral–dorsal) × 15 (rostral–caudal) mm^3^; matrix size = 256 × 128 × 48; spatial resolution = 59 × 59 × 313 µm^3^; 16 echoes per transient; and scan duration = 10.6 min.

DWI scans were performed in both 5xFAD and CADASIL models. Key imaging parameters were as follows:^31^ TR/TE = 2500/18.0 ms; field of view = 15 × 15 mm^2^; matrix size = 128 × 128; spatial resolution = 117 × 117 µm^2^; slice thickness = 0.75 mm; encoding direction = 12; *b* = 650 s/mm^2^; three repetitions of non-diffusion-weighted (*b* = 0 s/mm^2^) image; receiver bandwidth = 300 kHz; 16 axial slices; partial Fourier factor = 0.63 along the phase-encoding direction; and scan duration = 2.5 min with 4-segment SE EPI acquisition.

### Immunohistochemistry

Preparation and analysis of brain sections followed our previous reports.^31,32^ An iron stain kit (catalog no.: ab150674; Abcam, Cambridge, UK), also known as Prussian blue stain, was used to validate the cerebral microbleed.^33^ Staining procedures were performed in accordance with the manufacturer’s instructions.

### Data processing

All data processing was performed using custom MATLAB scripts (MathWorks, Natick, MA). GRE images (*in vivo* and *ex vivo*) were visually inspected by two raters, who reached consensus with discussion, to identify hypointense spots considered as microbleed when their sizes exceeded two pixels. SE images were used to estimate CSF volume and parenchymal tissue volume following a previously reported protocol. In addition, SE images were visually inspected for hyperintense spots outside the ventricular regions. DWI images were co-registered and normalized to a mouse brain template,^34^ and regions of interest (ROIs) encompassing the midbrain, cortex, hippocampus, thalamus, hypothalamus, and striatum were delineated. Regional ADC were quantified by spatial averaging the selected ROIs.

### Statistical analyses

When analyzing the 5xFAD data, an unpaired Student’s *t*-test was applied to examine group-wise differences in microbleed count (*in vivo* and *ex vivo*), parenchymal tissue volume, CSF volume, and regional ADC. Statistical details including *t*-statistics, degrees of freedom, and *p*-values were reported. The sex effect was evaluated using a linear regression model.

When analyzing the CADASIL data, linear mixed-effects models were used to examine the genotype (categorical factor), age (continuous factor), and genotype × age effects on water extraction fraction, PS, and regional ADC. A random intercept was included for mouse. Linear mixed-effects models were fit by restricted maximum likelihood (REML), with degrees of freedom estimated using the residual method. Effect estimate (denoted as β), 95% confidence interval (CI), and *p*-value were reported.

Data were presented as mean ± standard deviation. A *p*-value < 0.05 was considered statistically significant.

## Results

### Microbleed arose in 5xFAD but not in CADASIL mice

Representative GRE images of WT and 5xFAD mice illustrate the typical image quality obtained (Figure 2a). Microbleeds were observed in the hippocampus of both female and male 5xFAD mice (Figure 2a). At the group level, 5xFAD mice exhibited a higher microbleed count than WT controls (*t*[26] = 8.649, *p* < 0.001; Figure 2b). No sex-dependent difference was detected (linear regression: β = 0.246, 95% CI [–1.249, 1.741], *p* = 0.737). To validate the identification of microbleeds, Prussian blue staining was performed in representative mice. Positive staining was observed around the dentate gyrus region of the hippocampus in 5xFAD mice (Figure 2c), consistent with the MRI findings. No positive staining was found in matched WT controls (Figure 2c). As an additional control, brain sections from an intracerebral hemorrhagic stroke mouse model with intra-striatal collagenase injection^35^ showed positive staining surrounding the hemorrhagic area (Supporting Figure S1), confirming the validity of the staining procedure.

**Figure 2.**
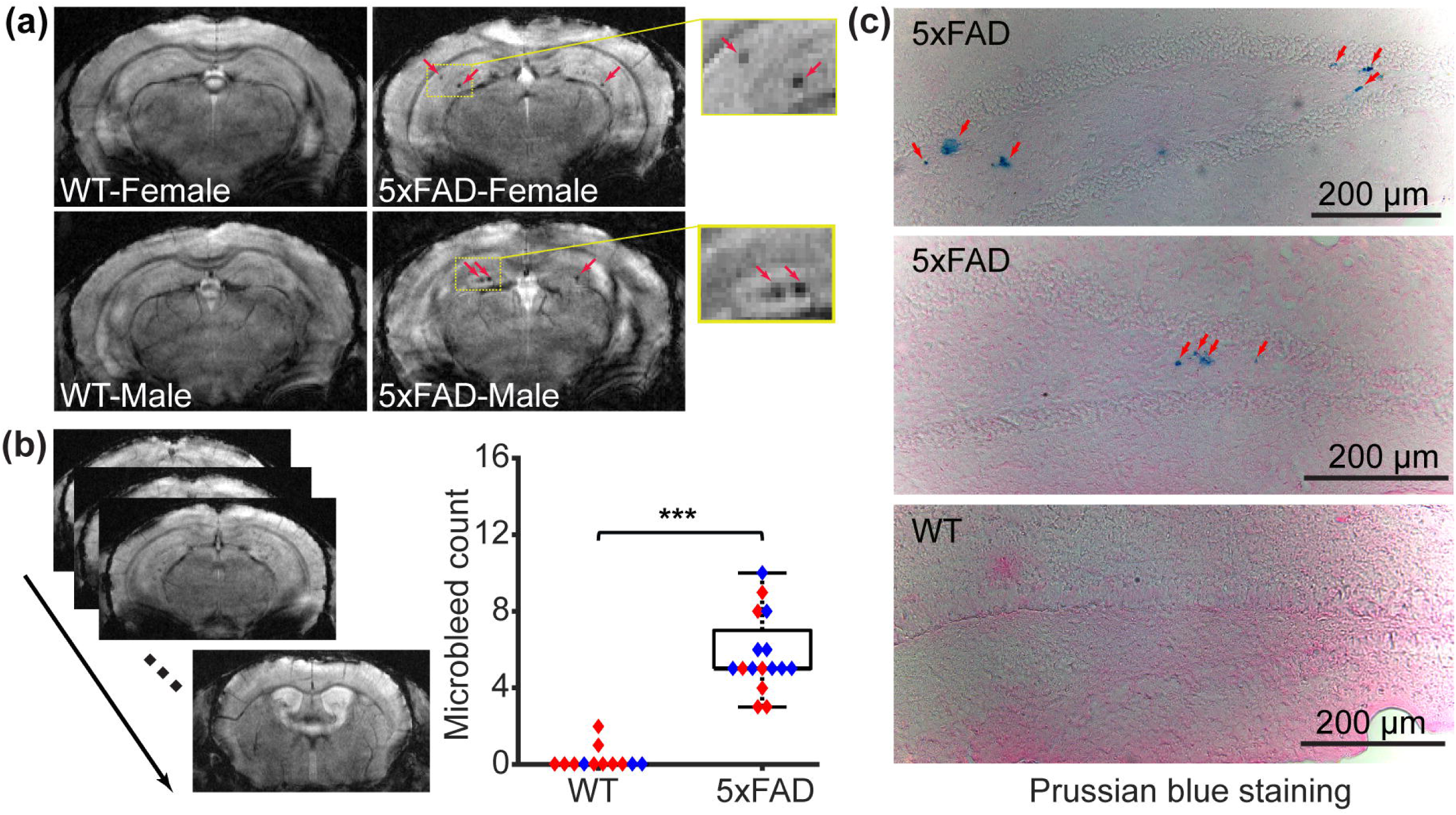
Microbleeds in the 5xFAD model (N = 28). (a) Representative gradient echo (GRE) images of female wildtype (WT), male WT, female 5xFAD, and male 5xFAD mice. (b) Slice-by-slice GRE images for a representative mouse and microbleed-count comparison between WT and 5xFAD mice. Red and blue dots represent female and male mice, respectively. In the boxplot, central mark was median, top and down edges of the box were 25^th^ and 75^th^ percentiles, and the whiskers extended to the minimal and maximal datapoints which were not considered to be outliers. \*\*\**p* < 0.001. (c) Microscope images with Prussian blue staining in 5xFAD and WT mice.

The *ex vivo* mouse brains were extracted and incubated in fomblin for imaging. Consequently, susceptibility-discontinuity-induced image distortion was reduced (Figure 3a). Slice-by-slice inspection confirmed good agreement in the identification of microhemorrhagic spots between *in vivo* and *ex vivo* datasets (Figure 3a). Owing to the enhanced through-plane resolution, microbleed spots were better captured in the *ex vivo* GRE images. The microbleed counts from *in vivo* and *ex vivo* GRE images were strongly correlated (Pearson’s correlation: R^2^ = 0.825, *p* < 0.001; Figure 3b). Referencing to the *ex vivo* data, 5xFAD mice exhibited significantly higher microbleed counts (*t*[8] = 4.704, *p* = 0.002; Figure 3c).

**Figure 3.**
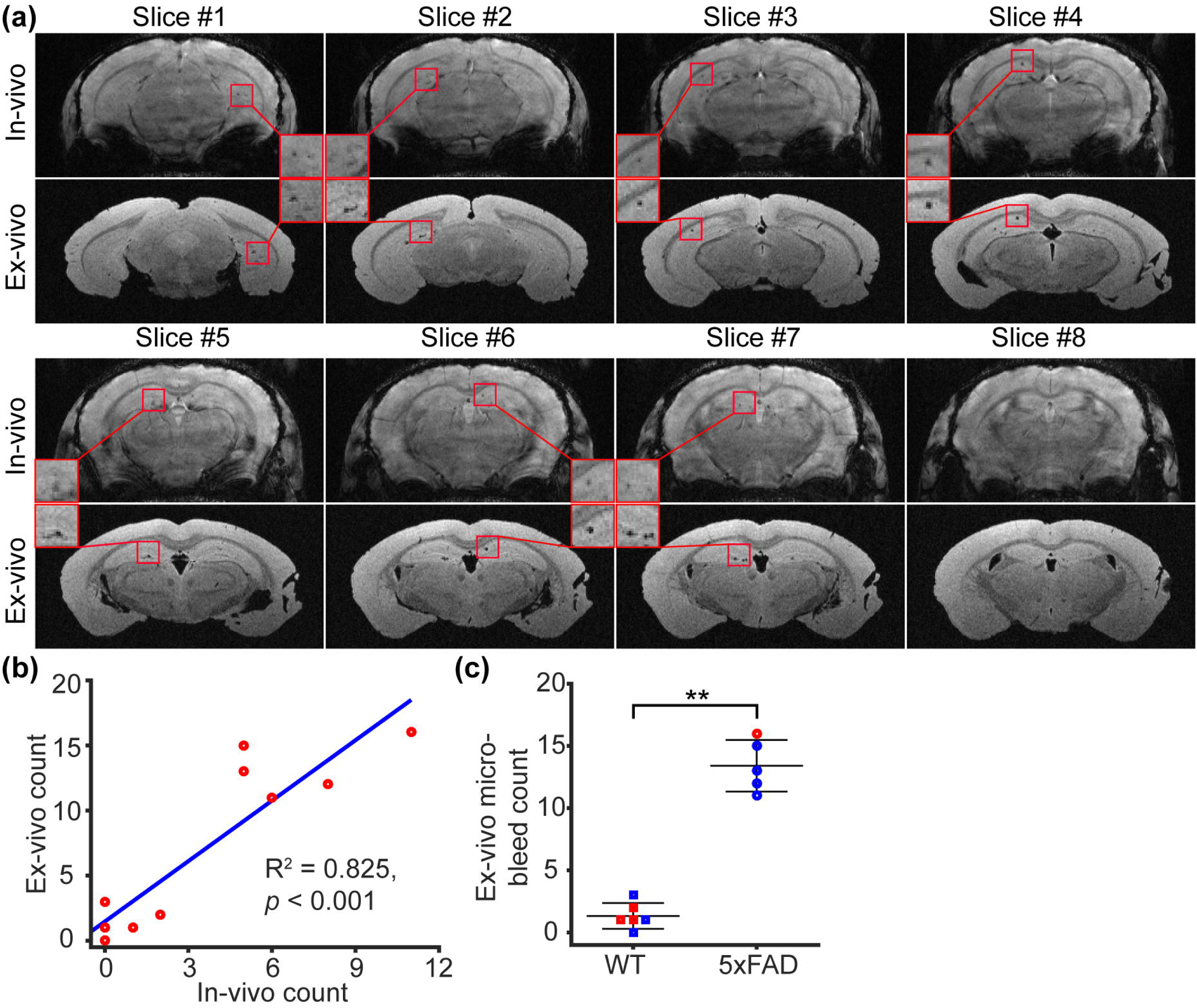
*Ex vivo* characterization of microbleed in the 5xFAD model (N = 10). (a) Comparison between *in vivo* and *ex vivo* GRE images in representative slices. (b) Correlation between *ex vivo* and *in vivo* microbleed counts. (c) Comparison of microbleed count between wildtype (WT) and 5xFAD mice using *ex vivo* MRI. Error bar represents standard deviation. \*\**p* < 0.01.

We examined the GRE images of WT and CADASIL mice at 9, 13, and 20 months of age and consistently found no hypointense spots. Representative images were displayed in Figure 4a. Consistent image quality was achieved for WEPCAST MRI across the three examined time points in the CADASIL model (Figure 4b), exhibiting reliable WEPCAST signals at the vein of Galen across all time points as marked by arrows (Figure 4b). Water extraction fraction exhibited an age effect (β = 0.07 %/month, 95% CI [0.01, 0.14], *p* = 0.028) but not a genotype (β = 0.15 %, 95% CI [-2.51, 2.81], *p* = 0.911) or sex effect (β = -0.08 %, 95% CI [-2.70, 2.53], *p* = 0.950) (Figure 4c). In a separate analysis for water extraction fraction, there was not a significant genotype × age effect (β = -0.02 %/month, 90% CI [-0.16, 0.11], *p* = 0.709). PS did not show a genotype (β = 7.16 ml/100g/min, 95% CI [-27.11, 41.43], *p* = 0.677), age (β = 0.59 ml/100g/min/month, 95% CI [-0.23, 1.41], *p* = 0.155), sex (β = 3.46 ml/100g/min, 95% CI [- 30.19, 37.11], *p* = 0.837), or genotype × age effect (β = 0.21 ml/100g/min/month, 95% CI [-1.46, 1.88], *p* = 0.802) (Figure 4d).

**Figure 4.**
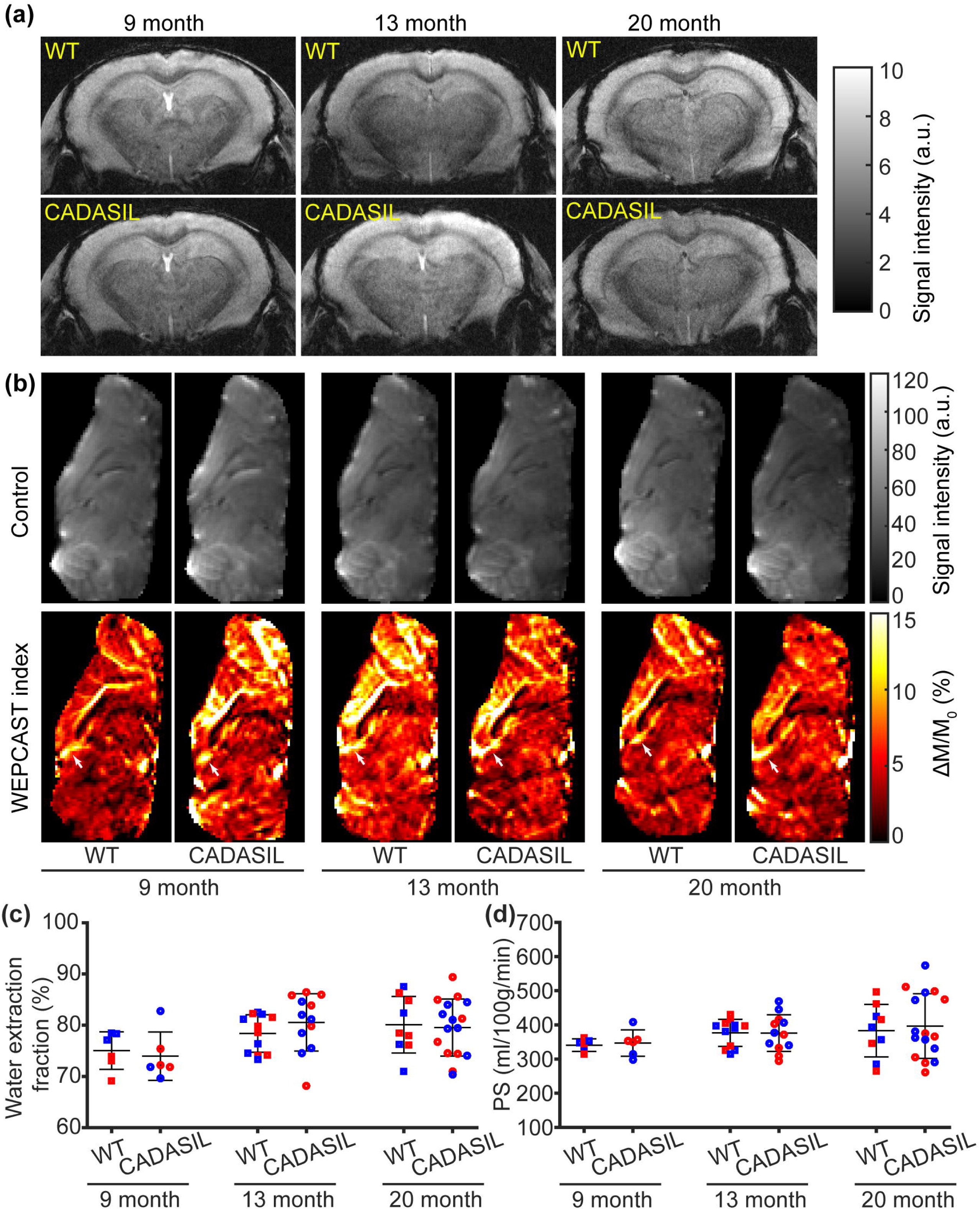
Gradient-echo (GRE) and water-extraction-with-phase-contrast-arterial-spin-tagging (WEPCAST) MRI of cerebral autosomal dominant arteriopathy and subcortical infarcts and leukoencephalopathy (CADASIL) model (N = 53). (a) GRE images for wildtype (WT) and CADASIL mice across the examined time points (9, 13, and 20 months of age). (b) Control and WEPCAST index images of WT and CADASIL mice across the time points. (c-d) Comparisons of water extraction fraction and permeability-surface area product (PS) across the examined time points for WT and CADASIL mice. Red and blue dots represent female and male mice, respectively. Error bar represents standard deviation.

### No hyperintensity or volumetric changes detected in 5xFAD mice (9–12 months)

No additional hyperintense spots were observed outside the ventricular regions in the SE images (Figure 5a-5b). At the group level, there were no significant differences in parenchymal tissue volume (*t*[26] = -0.512, *p* = 0.613; Figure 5c) or CSF volume (*t*[26] = 0.016, *p* = 0.988; Figure 5d), indicating the absence of genotype-dependent brain atrophy or ventricular enlargement. Separate linear regression analyses revealed no sex effect in parenchymal tissue volume (β = 1.135 mm^3^, 95% CI [-12.237, 14.508], *p* = 0.862) or CSF volume (β = -1.826 mm^3^, 95% CI [- 4.607, 0.955], *p* = 0.188).

**Figure 5.**
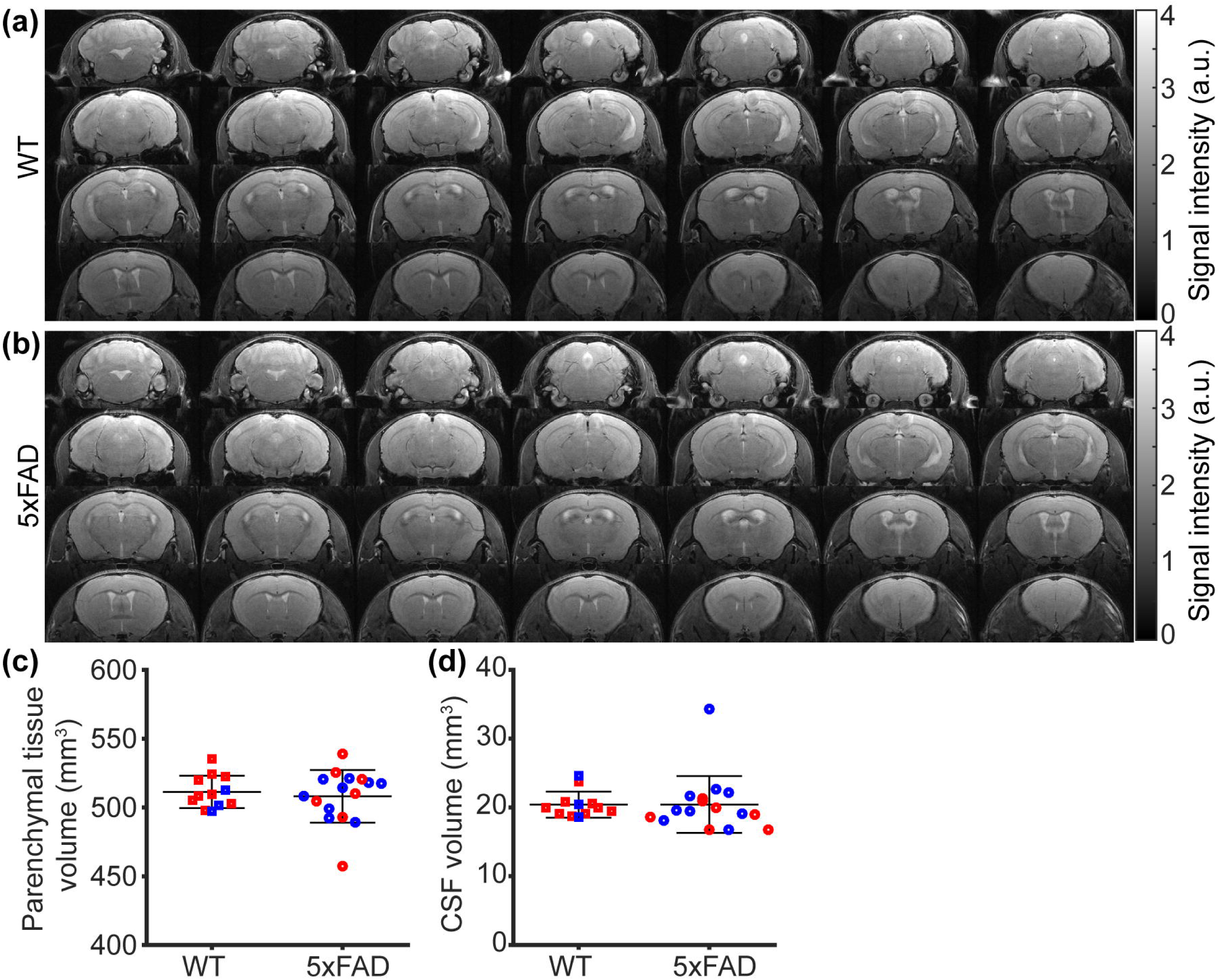
Spin echo (SE) MRI in the 5xFAD model (N = 28). (a-b) Representative SE dataset in wildtype (WT) and 5xFAD mice. (c-d) Comparisons of parenchymal tissue volume and cerebrospinal fluid (CSF) volume between WT and 5xFAD mice. Red and blue dots represent female and male mice, respectively. Error bar represents standard deviation. n.s.: not significant.

### Midbrain-specific ADC alteration in 5xFAD mice

Regional comparisons of ADC maps between WT and 5xFAD mice are shown in Figure 6a. A significant difference in ADC was detected in the midbrain (*t*[26] = 2.582, *p* = 0.016; Figure 6b), whereas no significant differences were observed in the isocortex (*t*[26] = 0.328, *p* = 0.745; Figure 6c), hippocampus (*t*[26] = –0.323, *p* = 0.750; Figure 6d), thalamus (*t*[26] = 0.332, *p* = 0.742; Figure 6e), hypothalamus (*t*[26] = 0.943, *p* = 0.354; Figure 6f), or striatum (*t*[26] = 0.615, *p* = 0.544; Figure 6g). Linear regression analyses revealed no significant sex effect (*p* ≥ 0.083) across the tested regions.

**Figure 6.**
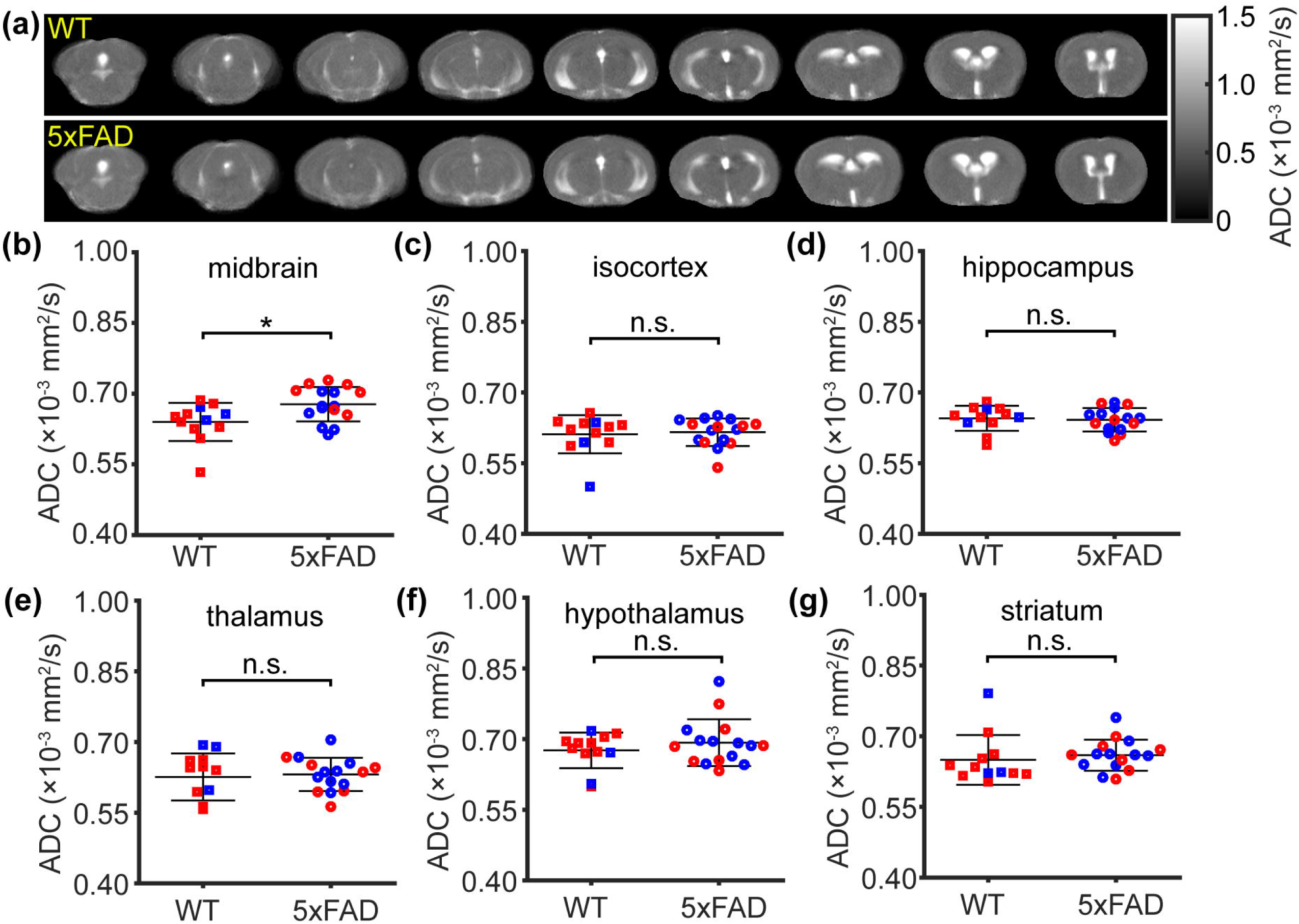
Regional apparent diffusion coefficient (ADC) in the 5xFAD model (N = 28). (a) mean ADC maps of WT and 5xFAD mice. (b–g) ADC comparisons between WT and 5xFAD mice in midbrain, isocortex, hippocampus, thalamus, hypothalamus, and striatum, respectively. Red and blue dots represent female and male mice, respectively. Error bar represents standard deviation. \**p* < 0.05. n.s.: not significant.

The CADASIL model exhibited a significant age effect on ADC across all tested brain regions (*p* ≤ 0.036) except for the isocortex (*p* = 0.085) (Supporting Figure S2 & Table S1). Such an age effect is consistent with findings from previous studies.^36,37^ No significant genotype effect, sex effect, or genotype × age interaction was observed (Supporting Figure S2 & Table S1).

The hippocampus exhibited microbleeds but normal ADC values, whereas the midbrain showed altered ADC without detectable microbleeds, suggesting a regional dissociation between microvascular rupture and diffusion-derived microstructural changes. Moreover, the observed ADC alterations were associated with amyloidosis rather than vascular smooth-muscle cell loss, indicating that amyloid-driven tissue pathology rather than vascular mural cell degeneration can lead to parenchymal microstructural abnormalities.

## Discussion

In this study, we systematically characterized microstructural alterations in mouse models of amyloidosis and vascular smooth-muscle cell loss. Microbleeds were associated with amyloid pathology in the absence of hyperintensity, brain atrophy, and ventricular enlargement, but not with vascular abnormalities arising from vascular smooth-muscle cell degeneration.

Our findings highlight the need to interpret cerebral microbleeds in a disease-specific framework and support the growing concept that cerebral microbleeds are a pathology-specific rather than a nonspecific SVD marker. In AD, microbleeds likely reflect a downstream consequence of CAA-related disruption of vascular integrity,^38^ serving as a potential imaging indicator of amyloid-driven neurovascular injury. In contrast, in non-amyloid SVD, where vascular smooth-muscle integrity, cerebrovascular reactivity, and baseline perfusion are markedly impaired, microbleeds are neither a consistent nor sensitive feature.^39,40^ This divergence underscores the potential of microbleeds as a disease-differentiating biomarker. Although microbleeds may not reliably indicate overall SVD severity, their spatial distribution and burden could provide valuable stratification for identifying AD patients with amyloid-mediated vascular fragility,^1,6^ thereby informing biomarker development and disease-specific therapeutic strategies.

Previous clinical and neuropathological studies have consistently shown that microbleeds in AD predominantly occur in lobar regions and are closely linked to CAA,^3,4^ whereas those in SVD typically localize to deep brain structures such as the basal ganglia, thalamus, and brainstem.^1^ This spatial distinction may reflect divergent underlying vascular pathologies. Our present findings reproduce these disease-specific patterns in experimental models, demonstrating that microbleeds emerge selectively under amyloid-driven vascular injury but not non-amyloid arteriopathy. This concordance across human and preclinical data supports the notion that microbleeds encode mechanistic information about the vascular substrate rather than serving as a generic marker of small-vessel injury.

BBB disruption and cerebral microbleeds represent two interrelated yet distinct manifestations of vascular dysfunction. BBB breakdown reflects increased permeability of the microvascular unit, allowing plasma proteins and other circulating components to enter the brain parenchyma,^41^ whereas microbleeds indicate structural rupture of small vessels leading to localized hemosiderin deposition.^1^ In AD, BBB leakage often precedes or accompanies amyloid-β accumulation in vessel walls, contributing to vascular inflammation, oxidative stress, and loss of mural cell integrity, thus enhancing the vulnerability of small vessels to rupture.^42^ Animal and human studies have consistently shown that regions exhibiting BBB impairment frequently co-localize with areas prone to microbleeds, suggesting a mechanistic continuum in which chronic barrier leakage weakens the vessel wall and eventually progresses to microbleed.^43^ Our previous study revealed a BBB disruption in 5xFAD mice starting from 3 months of age.^26^ In contrast, the present study demonstrates that CADASIL mice showed neither BBB permeability alterations nor microbleeds throughout the age range of 9 to 20 months. These findings are consistent with the proposed mechanistic continuum between BBB dysfunction and microbleed formation, suggesting that persistent barrier compromise in amyloid-bearing vessels may predispose them to later microvascular rupture, whereas non-amyloid arteriopathy without BBB leakage remains structurally stable. However, BBB disruption alone is not always sufficient to produce microbleeds; in non-amyloid SVD models, BBB permeability can be markedly elevated without detectable hemorrhagic foci.^44^ This dissociation implies that additional factors (e.g., amyloid deposition) are required for the transition from BBB dysfunction to frank microbleed formation. Collectively, BBB breakdown represents an early and potentially reversible stage of vascular compromise, while microbleeds may denote a more advanced and irreversible phase of vascular injury featuring the amyloid-laden vessels in AD.

AD develops through a complex cascade of interrelated pathological events^45^ involving amyloid accumulation, neuroinflammation, vascular dysfunction, metabolic dysfunction, and neurodegeneration. The process is thought to begin with amyloid-β overproduction or impaired clearance,^46^ leading to parenchymal plaque deposition and vascular amyloid accumulation.^3,4^ The prevalent amyloid-β deposition is accompanied by significant neuroinflammation.^21,47^ Vascular and metabolic dysfunctions are earlier events prior to significant brain atrophy.^48,49^ Within the context of these complications, microbleeds represent a downstream event of amyloid-driven vascular injury. Their presence on MRI provides a noninvasive window into the integrity of the neurovascular unit and serves as an imaging marker of advanced vascular pathology in AD. Recognizing microbleeds may serve not only as a diagnostic indicator of amyloid-related vascular compromise but also as a potential marker for monitoring disease progression and evaluating vascular-targeted therapeutic interventions. Considering microbleeds within the broader context of other pathological events in the AD cascade may further refine our understanding of the full landscape of AD pathophysiology.

ADC reflects the microstructural environment of water diffusion and primarily captures alterations in tissue integrity, such as cell swelling, edema, or neurodegeneration.^50^ In contrast, cerebral microbleeds represent focal vascular ruptures and hemosiderin deposition,^1^ which are primarily vascular phenomena rather than parenchymal microstructural changes. In our data, ADC alterations and microbleeds appeared as distinct, non-overlapping features, suggesting that microbleed formation does not necessarily coincide with detectable diffusion abnormalities at the tissue level. While ADC is sensitive to cytotoxic or degenerative processes within the parenchyma,^51^ microbleeds mark localized vascular damage without substantial diffusion perturbation in the surrounding tissue, highlighting that complementary information can be provided by integrating diffusion imaging with T_2_*-weighted imaging.

Our findings should be interpreted within the context of several limitations. First, although high-resolution gradient-echo MRI allowed sensitive detection of microbleeds, smaller lesions below the spatial resolution threshold may have been missed. Second, while our cross-sectional design captures representative stages of amyloid and vascular pathology, longitudinal tracking would better delineate the temporal relationship between amyloid deposition and microbleed formation in 5xFAD mice. Third, histological validation was limited to selected brain regions. Future studies incorporating spot-to-spot validation across multiple brain areas would further strengthen the findings. Despite these constraints, the present work provides a robust foundation for future translational efforts that integrate T *-weighted MRI with complementary molecular or perfusion biomarkers to refine vascular phenotyping in AD and related disorders.

## Conclusions

In summary, microbleeds hold potential as a disease-specific biomarker for detecting amyloid-related vascular injury and guiding the development of targeted diagnostic and therapeutic strategies in AD.

## Acknowledgments

This work was supported by the Grant Sponsors: National Institutes of Health (NIH) R01 AG081932 and NIH P41 EB031771. The authors thank Dr. Angeliki Louvi for advice on the generation and breeding of CADASIL model.

## Author contributions

X.Y.: Methodology, Investigation, Formal analysis, Visualization, Writing–Original draft, and Writing–Review & Edit. Y.L., A.B., and J.W.: Investigation and Writing–Review & Edit. M.W. and W.D.: Methodology, Resource, and Writing–Review & Edit. H.L.: Conceptualization, Methodology, Funding acquisition, and Writing–Review & Edit. Z.W.: Conceptualization, Methodology, Investigation, Software, Formal analysis, Visualization, Funding acquisition, Writing–Original draft, and Writing–Review & Edit.

## Supplemental material

Supplemental material for this article is available online.

