## Supplemental figures for "Vascular Microbleeds Without Brain Atrophy: A Microvascular Signature of Mid-Stage 5xFAD Pathology"

### Supplemental material

#### 1. Immunohistochemistry image of Prussian blue staining in an intracerebral hemorrhagic stroke model

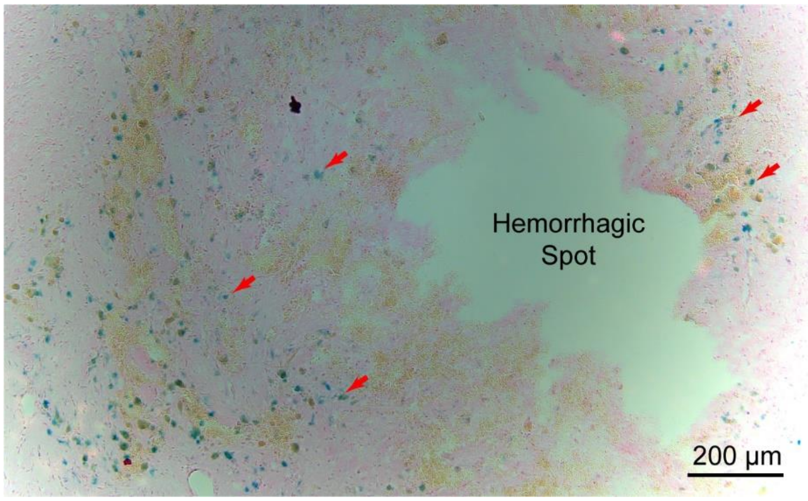


**Figure S1.** Prussian blue staining of a brain section from a mouse with intracerebral hemorrhagic stroke. The stroke model was generated by intra-striatal injection of collagenase. Red arrows indicate exemplary regions showing positive Prussian blue staining, corresponding to iron deposition.

### 2. DWI quantification in the CADASIL model

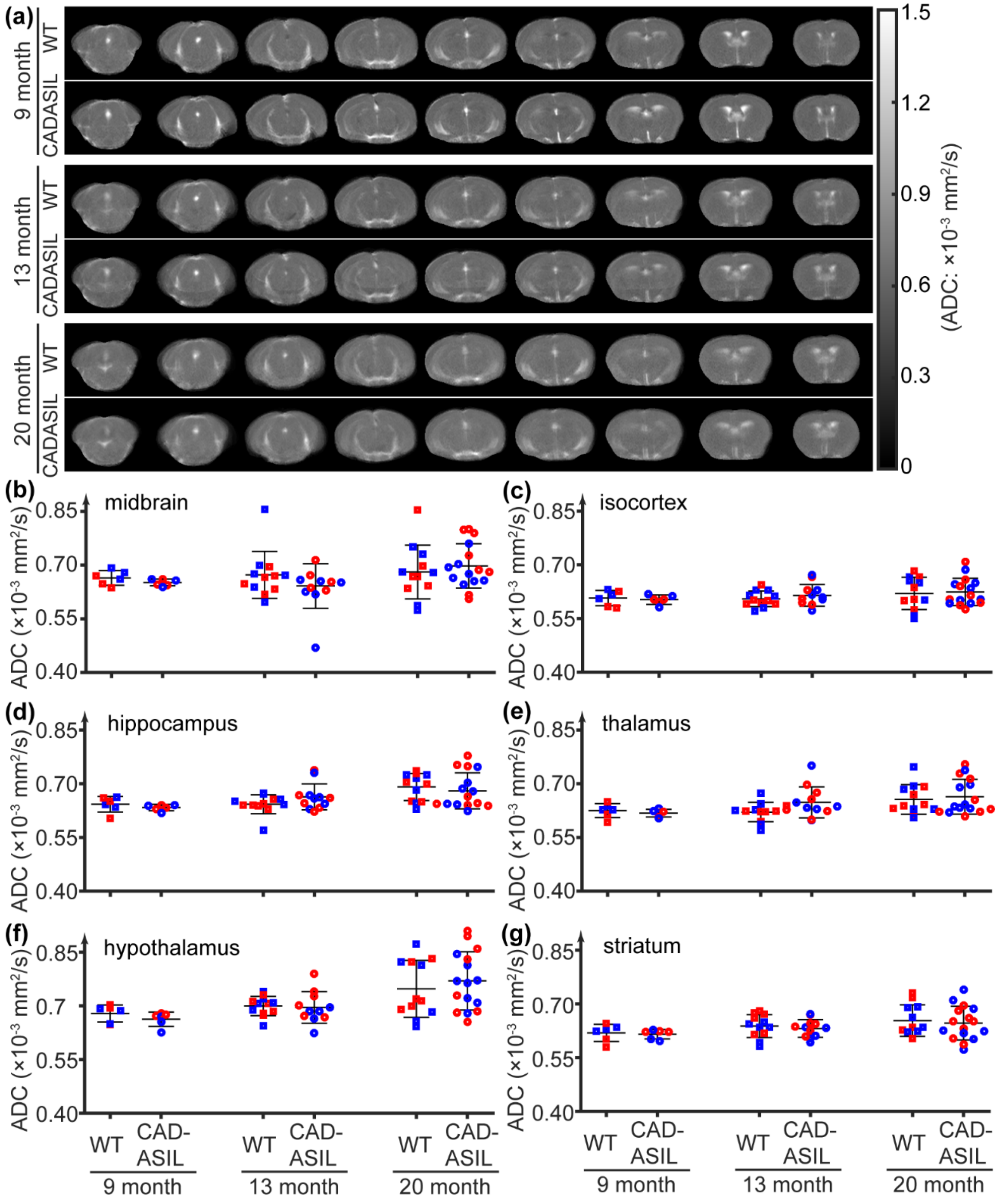


**Figure S2.** Averaged maps and regional quantification of ADC in the CADASIL model at 9, 13, and 20 months of age. (a) Averaged ADC maps. (b-g) Regional ADC across the tested time points for midbrain, isocortex, hippocampus, thalamus, hypothalamus, and striatum, respectively. Red and blue dots represent female and male mice, respectively.

**3. Statistical analyses of regional ADC values in the CADASIL model**

**Table S1** Detailed information of linear mixed-effects models

| Region | ADC ~ genotype + age + sex + (1\|mouse) | | | | | | | | | ADC ~ genotype × age + sex + (1\|mouse) | | |
|---|---|---|---|---|---|---|---|---|---|---|---|---|
| | genotype | | | age | | | sex | | | genotype × age | | |
| | β (×10$^{-5}$ mm$^2$/s) | 95% CI (×10$^{-5}$) | *p* | β (×10$^{-6}$ mm$^2$/s/month) | 95% CI (×10$^{-6}$) | *p* | β (×10$^{-5}$ mm$^2$/s) | 95% CI (×10$^{-5}$) | *p* | β (×10$^{-6}$ mm$^2$/s/month) | 95% CI (×10$^{-6}$) | *p* |
| midbrain | -0.65 | [-3.59, 2.28] | 0.657 | **3.58** | **[0.24, 6.94]** | **0.036** | 1.43 | [-1.49, 4.35] | 0.332 | 4.05 | [-2.58, 10.70] | 0.227 |
| isocortex | 0.49 | [-1.18, 2.15] | 0.562 | 1.49 | [-0.21, 3.19] | 0.085 | -0.21 | [-1.87, 1.45] | 0.799 | 0.16 | [-3.26, 3.57] | 0.927 |
| hippocampus | 0.23 | [-1.69, 2.16] | 0.808 | **3.85** | **[2.02, 5.69]** | **< 0.001** | 0.44 | [-1.48, 2.35] | 0.650 | -3.12 | [-6.51, 0.26] | 0.070 |
| thalamus | 1.35 | [-0.70, 3.41] | 0.193 | **2.80** | **[1.09, 4.51]** | **0.002** | -0.61 | [-2.65, 1.44] | 0.552 | -1.85 | [-5.07, 1.36] | 0.254 |
| hypothalamus | -1.83 | [-7.23, 3.58] | 0.501 | **7.89** | **[2.93, 12.84]** | **0.002** | 3.97 | [-1.42, 9.35] | 0.146 | 5.95 | [-3.68, 15.6] | 0.221 |
| striatum | -0.62 | [-2.43, 1.19] | 0.493 | **2.38** | **[0.51, 4.24]** | **0.014** | 0.58 | [-1.22, 2.39] | 0.521 | -0.27 | [-4.02, 3.48] | 0.888 |
